# Sex-specific survival but not tissue wasting in the KPP mouse model of pancreatic cancer-induced cachexia

**DOI:** 10.1101/2024.12.30.630679

**Authors:** Natalia M. Weinzierl, Jayarani Putri, Kathryn M. Spitler, Erin E. Talbert

## Abstract

Cancer cachexia, a multifactorial condition resulting in muscle and adipose tissue wasting, reduces the quality of life of many people with cancer. Despite decades of research, therapeutic options for cancer cachexia remain limited. Cachexia is highly prevalent in people with pancreatic ductal adenocarcinoma (PDAC), and many animal models of pancreatic cancer are used to understand mechanisms underlying cachexia. One such model is the *Kras*^*LSL-G12D*^, *Ptf1a*^*Cre-ER/+*^, *Pten*^*flox/flox*^ (KPP) model, which utilizes an inducible Cre recombinase to allow tumor development to be initiated at any age by tamoxifen administration. In our previous work, tumors were induced in KPP mice at 4 weeks of age. However, mice are still rapidly growing at this age, and a portion of the body weight differences seen between control and KPP mice is likely due to slowed growth of KPP mice. In our current study, pancreatic tumors were induced to develop with tamoxifen in KPP mice after rapid postnatal growth has slowed at 10 weeks of age (KPP10). Similar to our previous findings, KPP10 mice had lower body, muscle, and adipose tissue weights compared to non-tumor mice, and these differences were similar between male and female mice. However, male mice experienced greater relative weight loss. Unexpectedly, we identified that overall survival was significantly shorter in female KPP10 mice compared to KPP10 males. Greater body weight at tumor induction was associated with longer survival, suggesting that the sex difference in survival may be related to differences in body weight between male and female mice.

**NEW & NOTEWORTHY:**

- Although male mice experience greater relative body weight losses, similar skeletal muscle and adipose tissue wasting occurs between male and female mice in the *Kras*^*LSL-G12D*^, *Ptf1a*^*Cre-ER/+*^, *Pten*^*flox/flox*^ (KPP) model of pancreatic-cancer induced cachexia.
- Greater weight loss in males may be related to longer survival. However, differences in tamoxifen dose relative to body weight may have accelerated tumor formation in female mice and therefore may be a relevant consideration for inducible tumor models.

## INTRODUCTION

Weight loss, called cachexia, reduces the quality of life and outcomes for many people with cancer. Cachexia results from wasting of muscle and adipose tissue, affects 50-80% of all patients with cancer, and occurs more commonly in certain malignancies, such as pancreatic, gastric, and lung cancer (1). For example, nearly 80% of pancreatic cancer patients meet cachexia criteria at diagnosis (2, 3). Yet despite decades of clinical trials, no pharmacological therapies are approved to treat cancer cachexia outside of Japan (4).

The lack of an approved treatment may reflect differences in the underlying mechanisms of cachexia between patients and animal models used to develop anti-cachexia therapies. Many common models, such as the Colon-26 (C-26) and Lewis Lung Carcinoma (LLC) models, are characterized by a large tumor burden and short periods of rapid muscle wasting, which is not reflective of patients with cancer. Furthermore, high levels of circulating inflammatory cytokines appear to be critical for muscle wasting in these models, yet similar levels are rarely reported in cancer patients with cachexia (5-9).

Given these limitations, significant efforts have been undertaken to develop new mouse models of cancer cachexia, with emphasis on lung and pancreas cancers, as these two tumor types account for the largest number of patients with cachexia (1). For lung cancer, an orthotopic model has been developed (10) and an inhaled adenovirus tumor induction method has been repurposed for the study of cachexia (11, 12). In pancreas cancer, the common KPC mouse model (13) which utilizes a pancreatic progenitor cell-specific Cre recombinase (*Pdx1*^*Cre/+*^), a mutant form of Kras (*Kras*^*LSL-G12D*^), and either a mutant form (*Trp53*^*LSL-R172H*^ or *Trp53*^*LSL-R270H*^) or reduced expression (*Trp53*^*fl/+*^ or *Trp53*^*fl/fl*^) of the tumor suppressor p53 has been adapted to study cachexia (14, 15). Additional new pancreatic cancer cachexia models use intraperitoneal or orthotopic injections of cell lines derived from KPC tumors (16-19).

In a similar effort to study a model of pancreatic cancer-induced cachexia, we recently described the *Kras*^*LSL-G12D*^, *Ptf1a*^*Cre-ER/+*^, *Pten*^*flox/flox*^ (KPP) mouse (20). KPP mice develop pancreas tumors and exhibit both adipose tissue and skeletal muscle wasting. One strength of this model is that tumor development is inducible by tamoxifen, and therefore the age of tumor onset can be defined experimentally. In our previous work, we primarily initiated tumor development at 4 weeks of age and demonstrated a progressive loss of skeletal muscle and adipose tissue mass in KPP mice. However, since KPP mice are on a C57BL/6J background that continues to grow rapidly until 10 weeks of age (21), it is possible that a portion of the difference in tissue weights between KPP and control mice may have resulted from reduced growth, rather than tumor-induced tissue wasting.

To address this potential limitation in our mouse model, we sought to induce the development of pancreatic cancer at 10 weeks of age, when the rapid postnatal growth period of C57BL/6J mice has concluded. Since rates of muscle wasting have been shown to be higher in male versus female PDAC patients (14, 22), we were also interested in determining if the KPP model demonstrated similar differences in tissue wasting between male and female mice. Consistent with previous findings, KPP mice induced at 10 weeks of age (abbreviated KPP10) had lower body, muscle, and adipose tissue weights compared to non-tumor mice. Unexpectedly, we identified that relative weight loss was greater in male KPP10 mice and overall survival was significantly shorter in female KPP10 mice. Body weight at tumor induction positively correlated with survival, suggesting that survival among sexes may be related to differences in body weight between male and female mice. Taken together, our data point to advantages in using the KPP10 mice to study cancer cachexia, but also highlight considerations when using this model.

## MATERIALS AND METHODS

### Mice

KPP founder mice *Kras*^*LSL-G12D*^, *Pten*^*flox/flox*^ and *Ptf1a*^*Cre-ER/+*^, *Pten*^*flox/flox*^ were purchased from Jackson Labs (strain 033964). Mice were bred in house at The University of Iowa and were genotyped following The Jackson Laboratory protocol. These mice are maintained on a C57BL/6J background. Mice were weaned at 24 days of age, genotyped, and provided ad libitum access to diet and water. Mice were maintained on a 12/12 light cycle and housed with same-sex littermates. Control genotypes included 1) *Kras*^*LSL-G12D*^, *Pten*^*flox/flox*^; 2) *Ptf1a*^*Cre-ER/+*^, *Pten*^*flox/flox*^; and 3) *Pten*^*flox/flox*^.

At 10 weeks of age, a convenience sample of KPP10 and littermate control mice were injected intraperitoneally once daily for five consecutive days with 2 mg of tamoxifen in 100 µL of corn oil according to the protocol provided by The Jackson Laboratory (23). This induction time point was chosen because the rapid period of postnatal growth in C57BL/6J mice concludes shortly before 10 weeks of age in both male and female mice (21). Weight at the time of first injection was considered initial body weight.

All mice were weighed weekly to assess growth trends and identify the development of cachexia. Humane end-point criteria included weight loss of >20% of peak body weight, development of ascites, or failure to thrive. When endpoint criteria were reached, KPP10 mice and littermate controls were euthanized either by CO_2_ inhalation followed by cervical dislocation or by cervical dislocation under isoflurane anesthesia. A cohort of KPP10 mice and their littermate controls were weighed, euthanized under isoflurane anesthesia, and their muscle, adipose tissue, heart, liver, spleen, and pancreas including tumors were harvested.

Tissue weights are reported as wet weights. Muscle weights were averaged except in cases of obvious dissection errors. One KPP mouse was excluded for obvious liver disease unlikely to be related to the tumor pathology. When multiple mice were available, for example two control littermate mice for a single KPP10 mouse, a coin flip was performed to determine which mouse would be used as the comparator mouse. All experiments were approved by the University of Iowa Institutional Animal Care and Use Committee under protocols 9072241 and 2112241.

### Statistical Analysis

Peak body weight was defined as the highest weight achieved before consistent weight decline (i.e. cachexia onset). Low body weight was defined as the body weight at euthanasia following the draining of ascites. If tissues were not collected, low body weight was defined as the lowest body weight after the onset of cachexia, but before the development of ascites.

Body weight loss percentage was calculated as (peak body weight – low body weight) divided by peak body weight, then multiplied by 100 to yield a percentage. Differences between male and female KPP10 mice weight loss were assessed by a two-tailed unpaired t-test after an F-test was used to confirm equal variance between groups. Comparisons between KPP10 mice and controls for body and tissue weights were made by two-way ANOVA with sex and genotype (control/KPP) as the factors.

Survival days were counted from day 1 of tamoxifen administration until the day of euthanasia. Survival was compared via a Log-rank test. In addition to analyzing overall survival, we also investigated the time to peak weight, called cachexia latency.

The relationship between overall survival and pancreas + tumor weight, weight loss, body weight at the time of tumor induction, and peak body weight were analyzed via linear regression, with p values calculated by an F-test. All statistical tests were performed using GraphPad Prism 10. Statistical significance was defined as p<0.05. Data are reported as mean +/-standard deviation.

## RESULTS

### Sex-specific survival of KPP10 mice

After tumor induction at 10 weeks of age by five consecutive daily injections of tamoxifen, KPP10 mice and any littermates were euthanized when the KPP10 mouse reached any endpoint criteria, which included weight loss of >20% from peak body weight, development of visible ascites, or failure to thrive. KPP10 mice reached euthanasia criteria at a median of 136 days following tumor induction [Figure 1A]. KPP10 mice lost an average of 18.0 ± 9.0% of their peak body weight [Figure 1B]. Unexpectedly, female KPP10 mice had shorter survival compared to male KPP10 mice (median 124 days versus 149 days, respectively) [Figure 1C]. Male KPP10 mice lost significantly more weight (20.1 ± 9.8% of peak body weight) as compared to female KPP10 mice (15.4 ± 7.1%) [Figure 1D].

**Figure 1:**
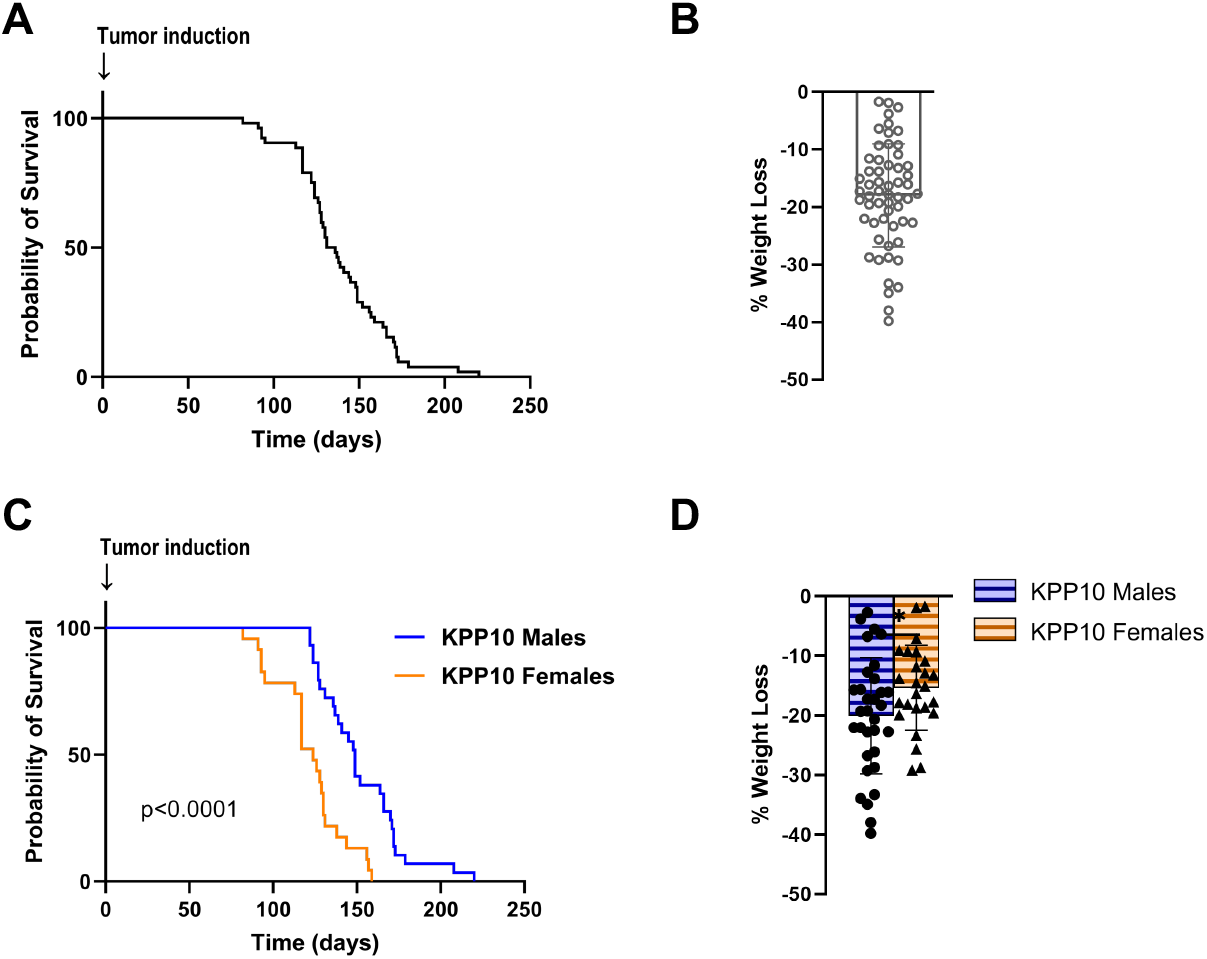
Survival and weight loss of KPP10 mice. [A] KPP10 mice had a median survival of 136 days from the first day of tumor induction (tamoxifen injection, 2 mg/day for 5 days, i.p.) until euthanasia. [B] Percent weight loss of KPP mice, calculated by (peak body weight minus ascites-low body weight) divided by peak body weight and multiplied by 100. [C] Male KPP10 mice had significantly longer survival than female KPP10 mice by a Log-rank test. Male KPP10 median survival = 149 days, female KPP10 median = 124 days. P<0.0001. [D] Male KPP10 mice had significantly greater weight loss than female KPP10 mice (20.1% ±9.8% versus 15.4%±7.1%, p=0.0489). *p<0.05 in an unpaired t-test. Total KPP10 mice = 57 [32 males, 25 females].

### Tumor burden and tissue wasting is similar between male and female KPP10 mice

To assess whether tissue wasting differed between male and female KPP10 mice, we compared a cohort of KPP10 mice to same-sex littermate controls. At euthanasia, KPP10 males and females had significantly lower ascites-free body weights compared to littermate controls [Figure 2A]. Consistent with our previous work (20), pancreas + tumor weights were significantly higher in KPP10 mice compared to controls [Figure 2B]. Neither body weight nor tumor weight demonstrated an interaction between sex and genotype, indicating that male and female KPP10 mice demonstrated similar differences in body weight and tumor burden when compared to control mice.

**Figure 2:**
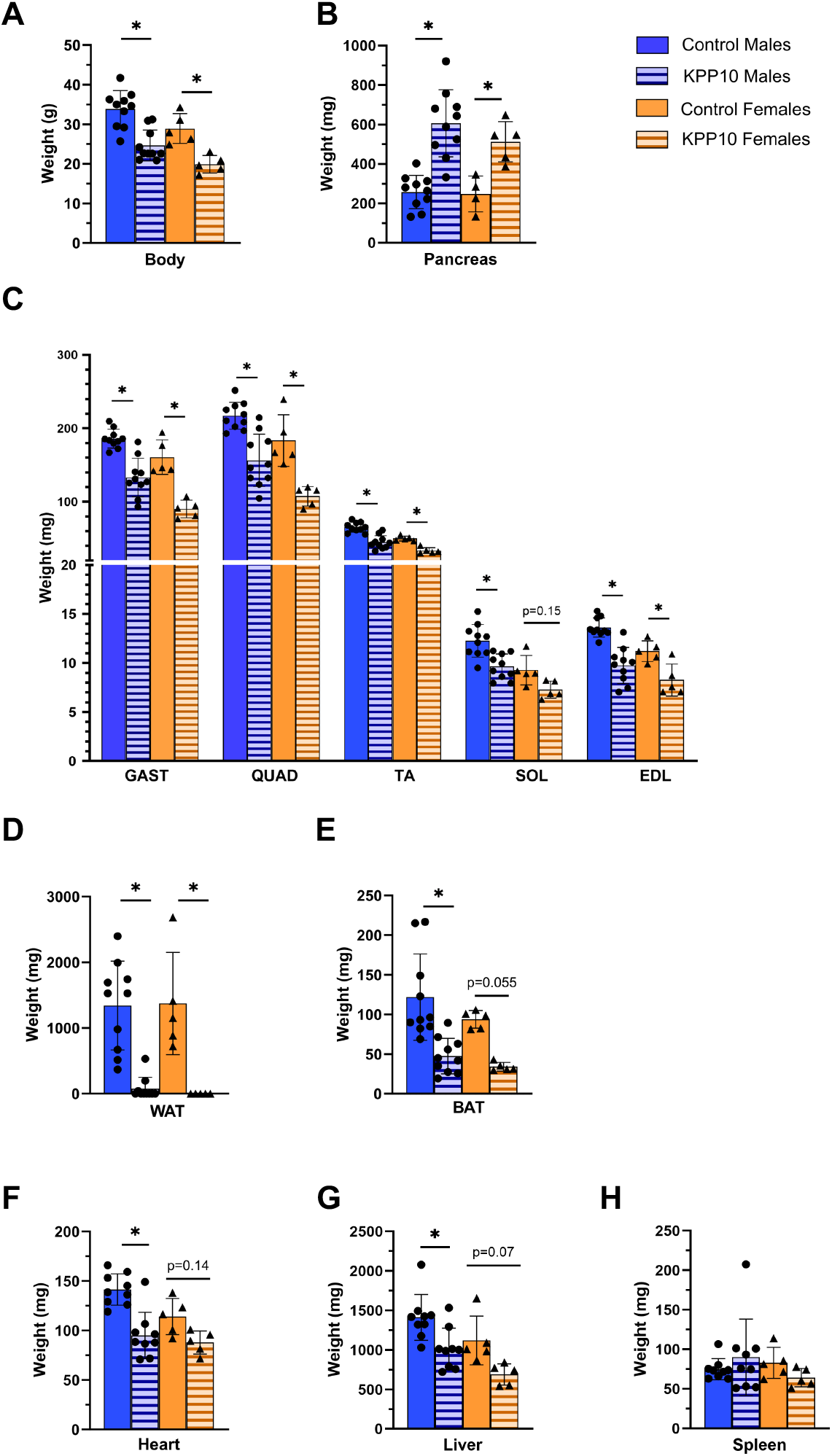
KPP10 mice exhibit reduced tissue masses. [A] KPP10 mice body weights are significantly lower than same-sex littermate controls (main effect for genotype, p <0.0001, main effect for sex, p=0.0039). [B] KPP10 mice have greater pancreas + tumor weights than control mice at euthanasia (main effect for genotype, p <0.0001). However, there was no difference between tumor + pancreas weights between male and female KPP10 mice (p=.31). [C] Gastrocnemius (GAST), quadriceps (QUAD), tibialis anterior (TA), soleus (SOL), and extensor digitorum longus (EDL) muscles all were smaller in KPP10 mice compared to controls (main effect for genotype: GAST p<0.0001, QUAD p<0.0001, TA p<0.0001, SOL p=0.0003, EDL p<0.0001; main effect for sex: GAST p=0.0002, QUAD p=0.0007, TA p<0.0001, SOL p<0.0001, EDL p= 0.0016). [D] KPP10 mice had less gonadal white (WAT) adipose tissue compared to controls (main effect for genotype, p <0.0001) [E] KPP10 mice had less subscapular brown adipose tissue (BAT) compared to controls (main effect for genotype, p<0.0001). [F] KPP10 mice had smaller heart weights at euthanasia compared to controls (main effect for genotype, p<0.0001, main effect for sex p=0.027). [G] KPP10 mice had smaller liver weights at euthanasia compared to controls (main effect for genotype, p=0.0005, main effect for sex p=0.007). [H] There were no differences in spleen weight between groups at euthanasia (genotype, p=0.88). n=10 males/group, n=5 females/group. Spleen, heart, and liver: n=9 males, n=5 females; pancreas n=10 males/group, n=4 control females, n=5 KPP10 females. Analyzed by two-way ANOVA with sex and genotype as factors. *p<0.05 in post-hoc analysis between indicated groups.

The masses of five hindlimb muscles (quadriceps, gastrocnemius, tibialis anterior, soleus, and extensor digitorum longus) were significantly smaller in KPP10 mice compared to littermate controls at euthanasia [Figure 2C]. KPP10 mice also had significantly less white adipose tissue (WAT) [Figure 2D] and brown adipose tissue (BAT) than controls [Figure 2E]. Heart weights were lower in KPP10 mice compared to control mice [Figure 2F]. In contrast to other cancer cachexia mouse models but consistent with our previous report, liver weights were significantly smaller in KPP10 mice compared to littermate controls [Figure 2G]. Spleen weights were not significantly different between KPP10 mice and controls [Figure 2H]. No interaction effects were present for any tissue, indicating a lack of sex differences in tissue wasting.

### Survival is associated with both tumor induction and peak body weight

Next, we sought to investigate potential causes of the shorter survival of female KPP10 mice [Figure 1C]. We found no association between pancreas + tumor weight and survival [Figure 3A, p=0.522] or percent weight loss and overall survival [Figure 3B, p=0.814]. We did, however, find a significant association between higher initial body weight and longer overall survival [Figure 3C, R^2^=0.161, p=0.002] and higher peak body weight and longer survival [Figure 3D, R^2^=0.286, p<0.0001]. To better characterize the survival difference between male and female KPP10 mice, we calculated cachexia latency, or the length of time between tumor induction and cachexia onset. We found that KPP10 females developed cachexia sooner (median=58 days) as compared to KPP10 males (median=111 days) [Figure 3E].

**Figure 3:**
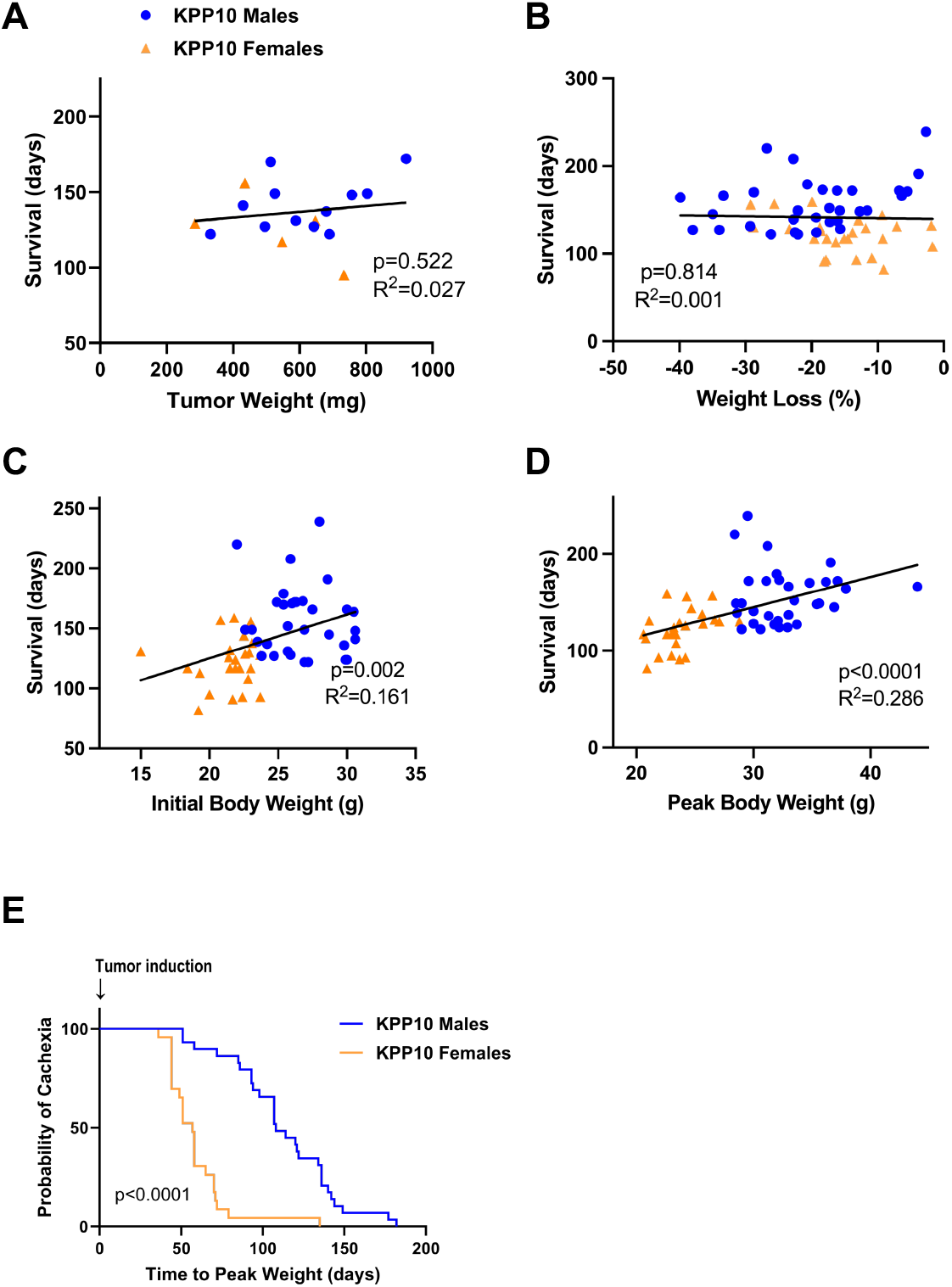
Sexual dimorphisms in cachexia latency and timeframe in KPP10 mice. No significant relationships existed between overall survival and pancreas + tumor weight (mg) at euthanasia [A] or weight loss (%) [B]. Higher initial body weight (g, at first tamoxifen injection) [C] and peak body weight (g) [D] were associated with increased overall survival. [E] Cachexia latency, defined as the time from induction to peak body weight, was longer in KPP10 males (111 days) than KPP10 females (58 days). [A]-[D] analyzed by simple linear regression; [E] analyzed by Log-rank test. [A]: n= 17 KPP mice, 12 males, 5 females. [B]-[E]: 57 KPP10 mice, 32 males, 25 females).

## DISCUSSION

Consistent with our previous findings (20), both male and female KPP10 mice exhibited significant losses in body weight and reduced adipose tissue and muscle masses compared littermate controls. KPP10 mice had an average of 31-33% lower masses of their quadriceps, gastrocnemius, and tibialis anterior muscles when compared to same-sex littermate controls [Figure 2C]. As expected, this difference was smaller than the 38-44% difference we reported in KPP mice induced at 4 weeks of age (KPP4), suggesting that reduced growth was likely a contributing factor to differences in muscle mass between control and tumor-bearing mice in our previous study (20). We recommend that users of the KPP model initiate PDAC induction no earlier than 10 weeks of age if mice are maintained on a C57BL/6J background. Furthermore, the need to delay tumor development until postnatal growth has concluded likely applies to any mouse model of cancer cachexia.

KPP10 mice had an overall survival of 136 days, which is longer than the 107 days that we previously reported for KPP4 mice (20). Unexpectedly, we identified a difference in median survival between female (124 days) and male (149 days) KPP10 mice [Figure 1C]. Sex differences in survival have not been previously reported in the KPC mouse model of pancreatic cancer (14, 24), although differences in cachexia development have been reported, with male mice developing cachexia more rapidly (14). Studies in humans with PDAC are mixed, with some studies suggesting females have modestly longer survival (25), with others finding no difference (26, 27).

We hypothesize that the survival difference between male and female KPP10 mice may not be inherent to the model, but instead is an artifact resulting from body weight at time of tamoxifen injection. All mice were injected with a standard dose of tamoxifen (2 mg/day for 5 days) per the protocol suggested by Jackson Labs (23). Because KPP10 females have a lower body weight than KPP10 males at 10 weeks of age, female mice received a higher dose of tamoxifen per body weight. Shorter survival [Figure 1C] and earlier cachexia onset [Figure 3E] in female KPP10 mice are consistent with faster tumor development, and therefore we hypothesize that KPP10 mice are sensitive to tamoxifen dosage. Anecdotally, female KPP10 mice also had a greater tendency to develop ascites compared to KPP10 males, which is consistent with increased tumor progression and early euthanasia in this group. Although the sample size was relatively small (n= 9 males, 3 females), a re-analysis of our previously published work showed a smaller, insignificant difference in survival in KPP4 mice (male median 110 days, female median 101 days, p=0.09). In this study, we adjusted tamoxifen dosages by weight, supporting our conclusion that not adjusting the dose of tamoxifen based upon mouse weight may have artificially induced the sex difference in survival. Of note, the KPC model does not require induction by tamoxifen for tumor development. In the KPC model, male mice have been reported to develop cachexia more rapidly (14), and this difference may help explain the differences between KPC mice and our findings in KPP10 mice.

We cannot exclude that exposure to endogenous estrogen in female KPP10 mice contributed to increased tumor development; however, we have previously reported that tamoxifen is required for activation of *Ptf1a*^*Cre-ER*^ (20). We believe that the higher weight loss seen in male KPP10 mice may be related to the longer survival of male mice [Figure 1C] and therefore may also be artifact of our tumor induction protocol.

Taken together, our data demonstrate that both male and female KPP10 mice experience cachexia. Further studies using weight-adjusted tamoxifen dosing will be required prior to undertaking studies to investigate sex differences using this model.

## DATA AVAILABILITY

The primary data are available on an Open Science Framework (OSF) Project site - DOI 10.17605/OSF.IO/275NP

## ACKNOWLEDGMENTS

The authors are grateful to Mikayla Kolpin, Lauryn Peck, Jacqueline Ott, Katherine Kavars, and Yabing Wang for critical feedback on the manuscript.

## AUTHOR CONTRIBUTIONS

NMW and EET conceived and designed the research, NMW, JP, KMS, and EET performed experiments, NMW and EET analyzed data, NMW and EET prepared figures, NMW and EET drafted the manuscript, and NMW, JP, KMS, and EET edited and approved final version of manuscript.

